# Dynamics of Bagaza, West Nile, and Usutu viruses in Portugal revealed by long-term serological surveillance in red-legged partridges, 2018-2022

**DOI:** 10.1101/2024.08.02.606292

**Authors:** Catarina Fontoura-Gonçalves, Francisco Llorente, Elisa Pérez-Ramírez, Miguel Ángel Jiménez-Clavero, João Basso Costa, Gonçalo de Mello, David Gonçalves, Paulo Célio Alves, Ursula Höfle, João Queirós

## Abstract

**Background:** Bagaza virus (BAGV) emerged in Portugal in 2021. BAGV and other flaviviruses such as West Nile virus (WNV) and Usutu virus (USUV) represent emerging threats in Europe, yet uncertainties persist regarding their epidemiological status in Portugal.

**Aim:** This study aimed to elucidate the epidemiological dynamics of BAGV, WNV, and USUV in Portugal using the red-legged partridge (*Alectoris rufa*) as a susceptible bird model species.

**Methods:** Serum samples collected from a partridge population in Southern Portugal, between 2018 and 2022, were analysed to characterize exposure to flaviviruses through enzyme-linked immunosorbent assay (ELISA; n=468) and virus neutralization test (VNT; n=433).

**Results:** ELISA test revealed an average seroprevalence of 58.1% for WNV and WNV-cross-reacting antibodies between 2018 and 2022, while 66.3% of individuals tested positive using VNT: 28.8% specific for WNV, 8.1% for BAGV, 2.1% for USUV and 27.2% for “undetermined flaviviruses”. BAGV seroprevalence was first detected in October 2021 (8.7%), alongside an outbreak detection, peaking in November 2021 (26.3%) and persisting until October 2022 (2.3%). WNV antibodies were detected throughout all sampling periods, with the highest seroprevalence in October 2020 (44.8%). USUV seroprevalence was detected until the BAGV outbreak in 2021, peaking in October 2020 (6.9%).

**Conclusion:** Long-term serological surveillance of partridges revealed endemic circulation of WNV and sporadic circulation of BAGV and USUV in Portugal. Notably, the highest WNV seroprevalence coincided with the human WNV outbreak in southern Spain in the summer of 2020, highlighting the role of red-legged partridges in the transmission/maintenance cycle and as sentinels of flaviviruses circulation.

## 1. Introduction

Emerging infectious diseases are a global health challenge, posing a threat to humankind and being responsible for serious morbidity and mortality events (Baker et al., 2021). West Nile virus (WNV), Usutu virus (USUV) and Bagaza virus (*Orthoflavivirus bagazaense,* BAGV) are positive-sense, single-stranded RNA viruses belonging to the mosquito-borne cluster of the *Orthoflavivirus* genus, family *Flaviviridae* (Benzarti et al., 2019). While BAGV recently emerged in Europe, WNV and USUV are considered re-emerging viruses in Europe, with zoonotic potential and associated with neurological disease (Baker et al., 2021). These flaviviruses are maintained in nature mainly through an enzootic life cycle involving mosquitoes as vectors and birds as the main reservoir host (Benzarti et al., 2019). Incidental infection of other hosts, such as humans and horses, can occur, but they are considered dead-end hosts without the ability to transmit the pathogen (Benzarti et al., 2019). In the last decade, the increased epidemic activity accompanied by an upsurge in human morbidity and mortality has placed avian flaviviruses as a public health concern (Rodríguez-Alarcón et al., 2021).

BAGV belongs to the Ntaya serocomplex and was first detected in the Bagaza district of Central African Republic in 1966 in a pool of *Cule*x mosquitos (Digoutte, 1978). In 2010 the virus was detected for the first time in Europe in an outbreak in Southern Spain (Llorente et al., 2013) that affected red-legged partridges (*Alectoris rufa*) and pheasants (*Phasianus colchicus*). Later it was found to be equivalent to Israel turkey meningoencephalitis virus (Fernández-Pinero et al., 2014), already associated with a serious disease affecting poultry (turkeys). Since 2010, BAGV has been detected sporadically in Southern Spain, usually in cocirculation with WNV and/or USUV (Llorente et al., 2013; Varga et al., 2024). Recently, BAGV also caused a disease outbreak in the South of Portugal (Queirós et al., 2022) and Spain (Höfle et al., 2022), affecting mainly partridges, but also a songbird, the corn bunting (*Emberiza calandra*) (Queirós et al., 2022). This has raised the question of whether BAGV was introduced in Portugal for the first time in 2021 or if it had been silently circulating in previous years, with undetected mortality. In the 2010 outbreak in Spain, the virus caused a mortality rate of 23% in partridges based on epidemiological enquiries (García-Bocanegra et al., 2013), whilst experimental studies in the same species have reported a 30% mortality rate (Llorente et al., 2015).

WNV and USUV belong to the Japanese Encephalitis virus (JEV) serocomplex (Benzarti et al., 2019). Both viruses co-circulate in many regions (Nikolay, 2015) with increasing incidence and distribution in Europe (Vilibic-Cavlek et al., 2020). Regarding WNV, the virus has been circulating in Spain since at least 2005. However, in 2020 it caused the largest epidemic so far, with 77 human cases and 8 deaths (Figuerola et al., 2022; Rodríguez-Alarcón et al., 2021). In Portugal, the endemic circulation of WNV remains unclear, with only four serologically confirmed human infections in the last 40 years, and few studies addressing the epidemiological situation in vertebrates and vectors (a few from 2004-2010 in birds, mosquitos and horses and more recently some additional reports in equines mainly in Alentejo and Algarve) (Lourenço et al., 2022).

USUV circulation is known in Europe since 1996 (Weissenböck et al., 2013), causing a large epidemic in birds (blackbirds, *Turdus merula*) in 2001 in Austria, from where it spread to central Europe causing sporadic outbreaks in birds (Vilibic-Cavlek et al., 2020). It has been detected in several species across Europe (birds, mammals, mosquitoes), and seems to be highly pathogenic for some bird species, namely blackbirds. Its zoonotic potential has also been confirmed with human cases reported in eight European countries (Stephan et al., 2018). In Portugal, however, no human or animal cases of USUV have been identified. In Spain, the virus was detected for the first time in mosquitos in 2006 (Busquets et al., 2008), and since then, it has been reported in wild birds (Höfle et al., 2013) and horses (Magallanes et al., 2023).

The red-legged partridge is a game species of significant ecological, economic and social importance in Portugal and Spain (Farfán, 2022). It is susceptible to flaviviruses infection and may act as a competent host for amplification of WNV (Pérez-Ramírez et al., 2018).

In this study, we aimed to understand the epidemiological dynamics of WNV, USUV and BAGV in wild birds in Portugal, with particular focus in BAGV as it caused an important outbreak in 2021 (Queirós et al. 2022). For this, we used the red-legged partridge as a susceptible bird model species for flaviviruses (Pérez-Ramírez et al., 2018) and a long-standing surveillance program implemented in Southern Portugal.

## 2. Material and methods

### 2.1 Study area, sampling, and data collection

This study was conducted in a hunting estate in the Municipality of Serpa (37.82236N, - 7.37953W), Southern Portugal, between October 2018 and October 2022 (Figure 1).

**Figure 1.**
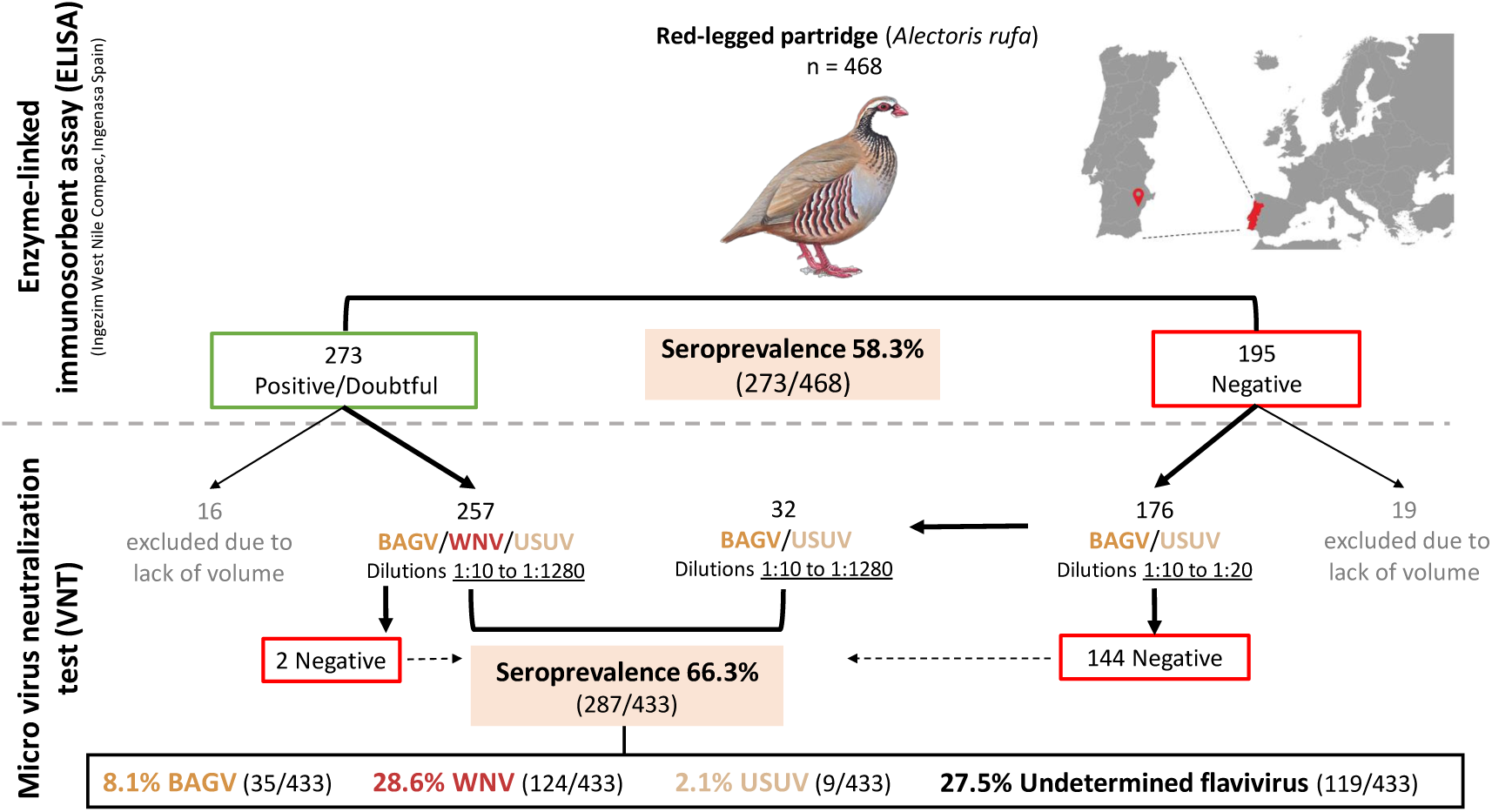
Location of the red-legged partridge population sampled in this study and the diagnostic protocol used to quantify the seroprevalence of WNV and WNV cross-reacting antibodies by ELISA and specific seroprevalence for BAGV, WNV, USUV and “undetermined flavivirus” by VNT.

During these five years, partridges were sampled at least once a year, after the reproductive season (October-December). In 2019, an additional sampling was performed in February. After the BAGV outbreak in 2021 (Queirós et al., 2022), we increased the sampling effort and samples were collected approximately every two months (Table S1 indicates the sampling time points and number of samples per sampling event). Samples were obtained either after the live capture of partridges or during necropsies of hunted/dead specimens. Live partridges were captured using baited walk-in traps. Each partridge was ringed and age and sex recorded. The age was determined by examining the primary feathers, while the sex was identified based on spur characteristics. Blood (<1% body mass) was collected using 1 ml syringes and stored in sterile eppendorf tubes. Partridges were released at the capture site after sampling. In total, we collected 468 blood samples that were allowed to cloth at room temperature and then maintained at 4°C for at least 2 h and a maximum of 24 h before centrifugation for 10 min at 12,000 rpm to separate serum and cellular fractions. Serum samples were maintained at -20°C until further analysis. Legal permissions to capture and mark the animals were provided annually by the Portuguese National Institute of Wildlife Conservation and Protection (ICNF).

### 2.2 Antibody detection assay

Initial screening for flavivirus antibodies was performed using a commercial competitive ELISA (Ingezim West Nile Compac, Ingenasa Spain), following the manufactureŕs instructions. Despite being designed for the detection of WNV antibodies, this diagnostic test may cross-react with antibodies directed against other flaviviruses, like USUV and BAGV (Llorente et al., 2019).

To assess the specific seroprevalence of each virus a micro virus neutralization test (VNT) against WNV, USUV and BAGV was performed, as it is considered the gold standard method for WNV serological diagnosis (OIE, 2013). WNV lineage 1 strain Eg-101/NY99, USUV strain SAAR1776 and BAGV strain Spain/2010 were used for the VNT in Vero cell cultures as previously described (Llorente et al., 2019). The VNTs were performed in the BSL-3 laboratory at *Centro de Investigación en Sanidad Animal* (CISA-INIA), CSIC. In the case of positive and doubtful ELISA samples, neutralizing antibody titres were determined in parallel for each serum sample against the three viruses using serial (twofold) serum dilutions (1:10-1:1280/10240), while negative ELISA samples were analysed in parallel only against USUV and BAGV using two serial dilutions (1:10-1:20), due to the higher specificity of the ELISA test to WNV when compared with USUV and BAGV (Llorente et al., 2019). In this latter case, samples with no neutralizing immune response (<1:10) were considered negative for the three viruses, whereas all others were considered positive and again tested by VNT to USUV and BAGV in parallel, this time using serial dilutions 1:10-1:1280 (Figure 1).

Specific neutralizing antibody responses were based on the comparison of VNT titres obtained against the two/three flaviviruses: the neutralizing immune response observed was considered specific for a given virus when VNT titres were at least fourfold higher than the titre obtained against the other viruses. When VNT titre differences did not reach this threshold, the result was considered inconclusive and the specificity for a given flavivirus could not be determined (“undertmined flaviviruses”; Llorente et al., 2019). For 35 samples VNT could not be performed due to insufficient serum volume.

### 2.3 Individual and temporal drivers of flavivirus exposure

To determine the effect of individual (sex and age class) and temporal (year and month) drivers of flavivirus exposure in red-legged partridges, general linear models (GLM) with binomial distribution and logit link function were constructed. As dependent variable, we used the individual serological status, classified by the results of ELISA and VNT: WNV, BAGV, USUV, “undetermined flaviviruses” and general VNT (all the positives to VNT). Since the sampling effort was not equal in all years, two data sets were analysed: i) considering just one sampling per year, in autumn; and ii) considering the samplings every two months since the BAGV outbreak in September 2021. Distinct GLMs were constructed for each of the previous data sets. All statistical analyses were performed using R software, version 4.3.2.

## 3. Results

ELISA results revealed an overall WNV and/or cross-reacting flavivirus seroprevalence of 58.3% (273/468) in partridges sampled between 2018 and 2022 (Figure 1; Table S1). In contrast, VNT results (Table S2) revealed an overall flavivirus seroprevalence of 66.3%, classified as WNV (28.6%, 124/433), USUV (2.1%, 9/433), BAGV (8.1%, 35/433) or “undetermined flaviviruses” (27.5%, 119/433) (Figure 1). The discrepancy between ELISA and VNT resided in two positive ELISA samples that were classified as negative by VNT and in 32 ELISA negative samples, of which 19 were positive to BAGV, one to USUV and 12 were classified as “undetermined flaviviruses”.

VNT seroprevalence to the different flavivirus varied between autumns (Figure 2). BAGV antibodies were first detected in the autumn of 2021 with a seroprevalence rate of 16.3% (14/86) and decreased to 2.3% (1/44) in 2022 with only one positive individual detected. WNV seroprevalence ranged from 24.0% (12/50) in 2018 to 44.8% (13/29) in 2020 when the highest rate was detected, then decreasing to 26.7% (23/86) in 2021 and 36.4% (16/44) in 2022. For USUV, in the autumn of 2018 a seroprevalence of 6.0% (3/50) was observed.Specific antibodies were again detected in 2020 and 2021 with seroprevalence rates of 6.9% (2/29) and 3.5% (3/86), respectively. USUV antibodies were not identified in 2019 (0/39) nor in 2022 (0/44). The seroprevalence rate to “undetermined flaviviruses” was 22.0% (11/50) in 2018, stayed roughly similar in 2019 with 20.5% (8/39) and in 2020 with 20.7% (6/29), but peaked in 2021 and 2022 with 33.7% (29/86) and 31.8% (14/44), respectively.

**Figure 2.**
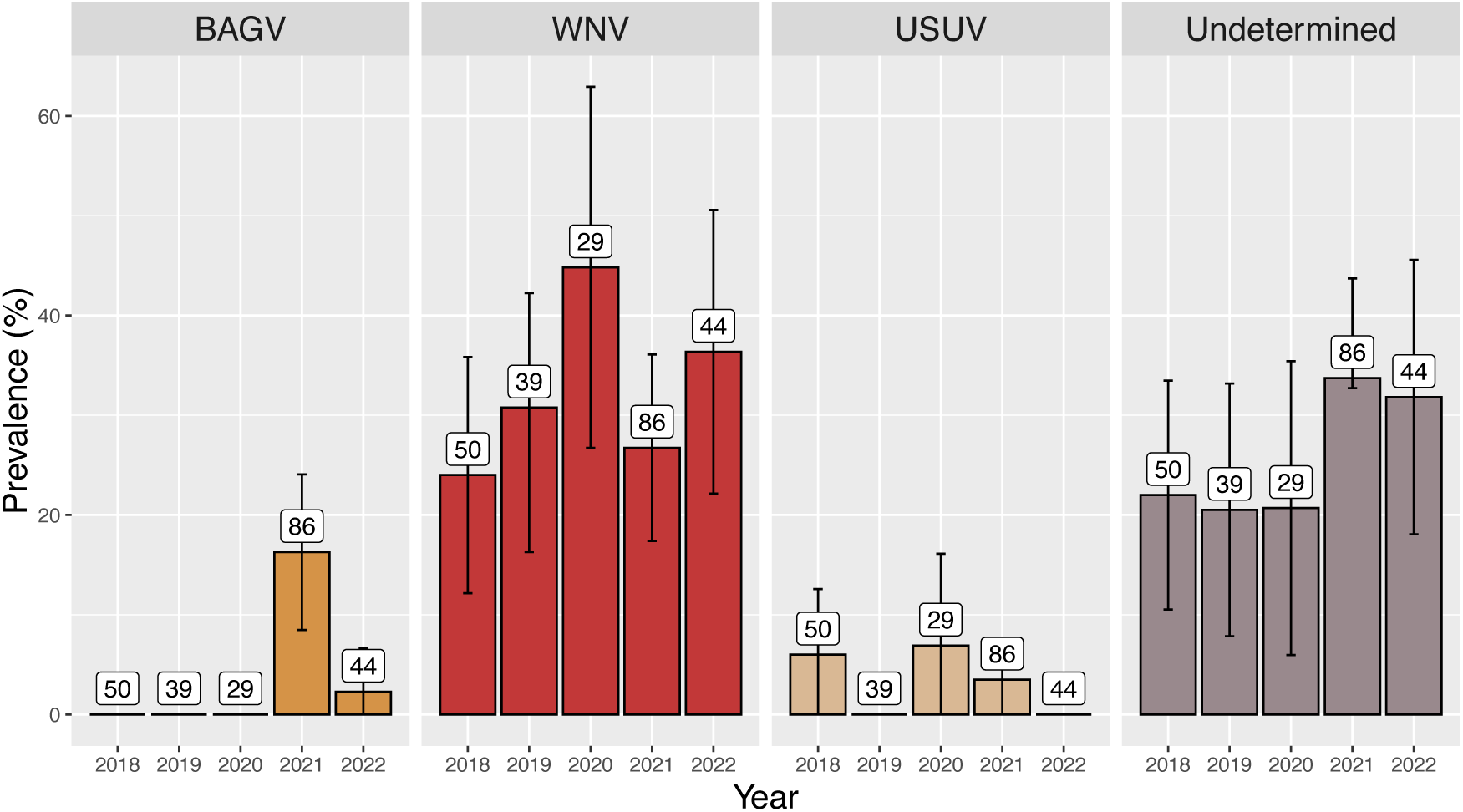
Seroprevalences obtained from the VNT for BAGV, WNV, USUV and Undetermined (“undetermined flaviviruses”) in the autumn of all years. The total number of samples analysed is indicated in each column and error bars depict the 95% confidence interval, calculated assuming a normal distribution.

VNT seroprevalence to the different flaviviruses also varied between the bi-monthly sampling time points (after the BAGV outbreak was detected) comprised between September/October 2021 to October 2022 (Figure 3). BAGV specific antibodies were first detected in September/October 2021 with a seroprevalence of 8.3% (4/48). In November 2021, the highest seroprevalence was detected, 26.3% (10/38) after which it decreased until October 2022 with 2.3% (1/44). WNV antibodies were detected in all sampling periods, with the lowest seroprevalence of 9.5% (4/42) detected in August 2022 and the highest seroprevalence of 44.8% (13/29) in October 2020. USUV antibodies were detected in September/October 2021 with a seroprevalence of 4.2% (2/48) and in November 2021 with 2.6% (1/38) and from there USUV antibodies were no longer detected.

**Figure 3.**
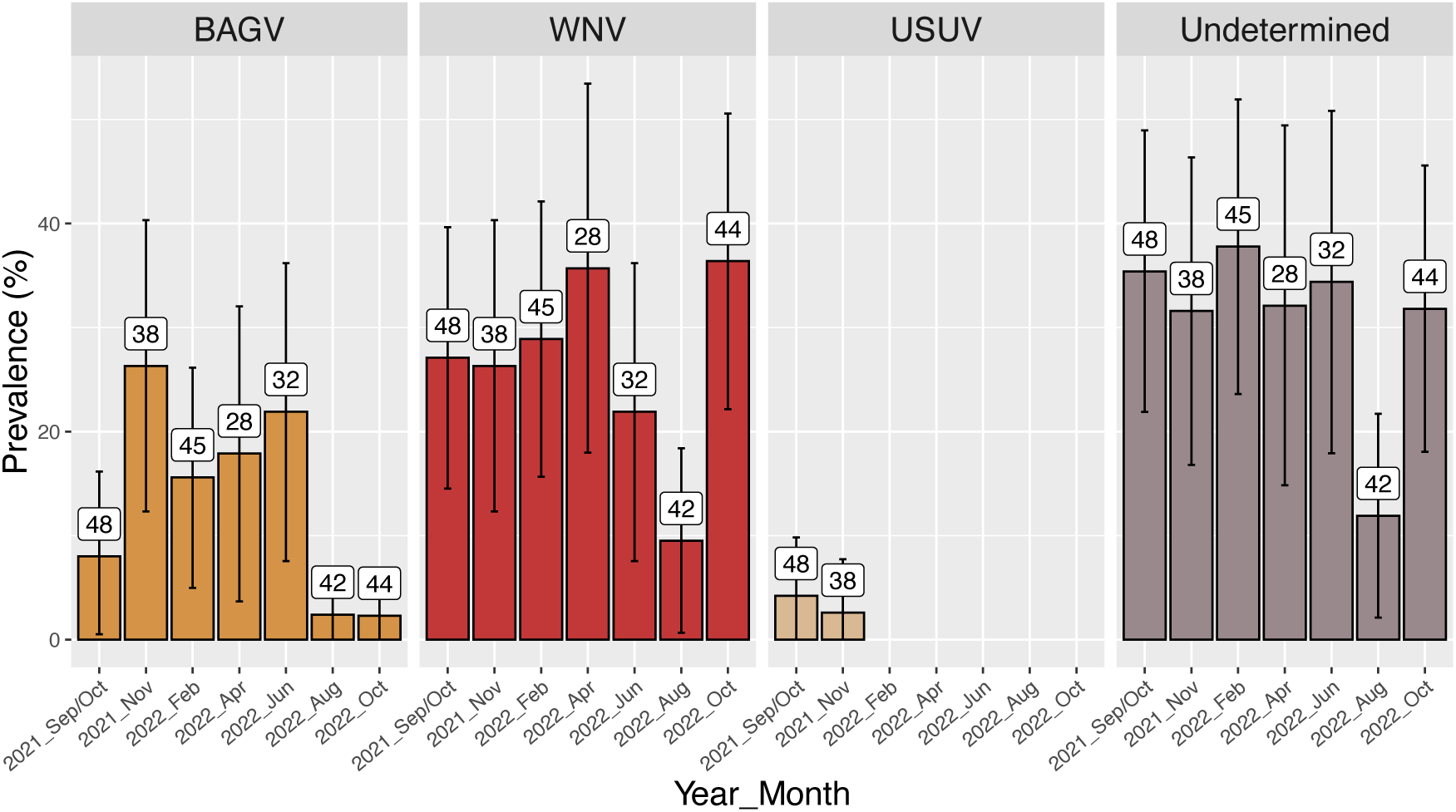
Seroprevalences, obtained by VNT, for BAGV, WNV, USUV and Undetermined (“undetermined flaviviruses”) at two-month sampling intervals after the BAGV outbreak. The total number of samples analysed is indicated in each column and error bars depict the 95% confidence interval, calculated assuming a normal distribution.

Antibodies against “undetermined flaviviruses” were present throughout all sampling periods, with the lowest seroprevalence of 11.9% (5/42) found in August 2022 and the highest with 37.8% (17/45) in February 2022 (Figure 2). The most prevalent undifferentiable titres against more than one virus were between the WNV/USUV and WNV/USUV/BAGV (Figure S1). Neutralizing flavivirus antibody titres ranged from 1:10 to >1:10240 (Figure 4, Table S3). The highest individual titres were detected for individuals sampled in September/October and November 2021, both for WNV, USUV and BAGV (Figure 4, Table S3). The highest mean titres for WNV and BAGV were also detected in September/October 2021 and for USUV in November 2021 (Figure 4, Table S3).

**Figure 4.**
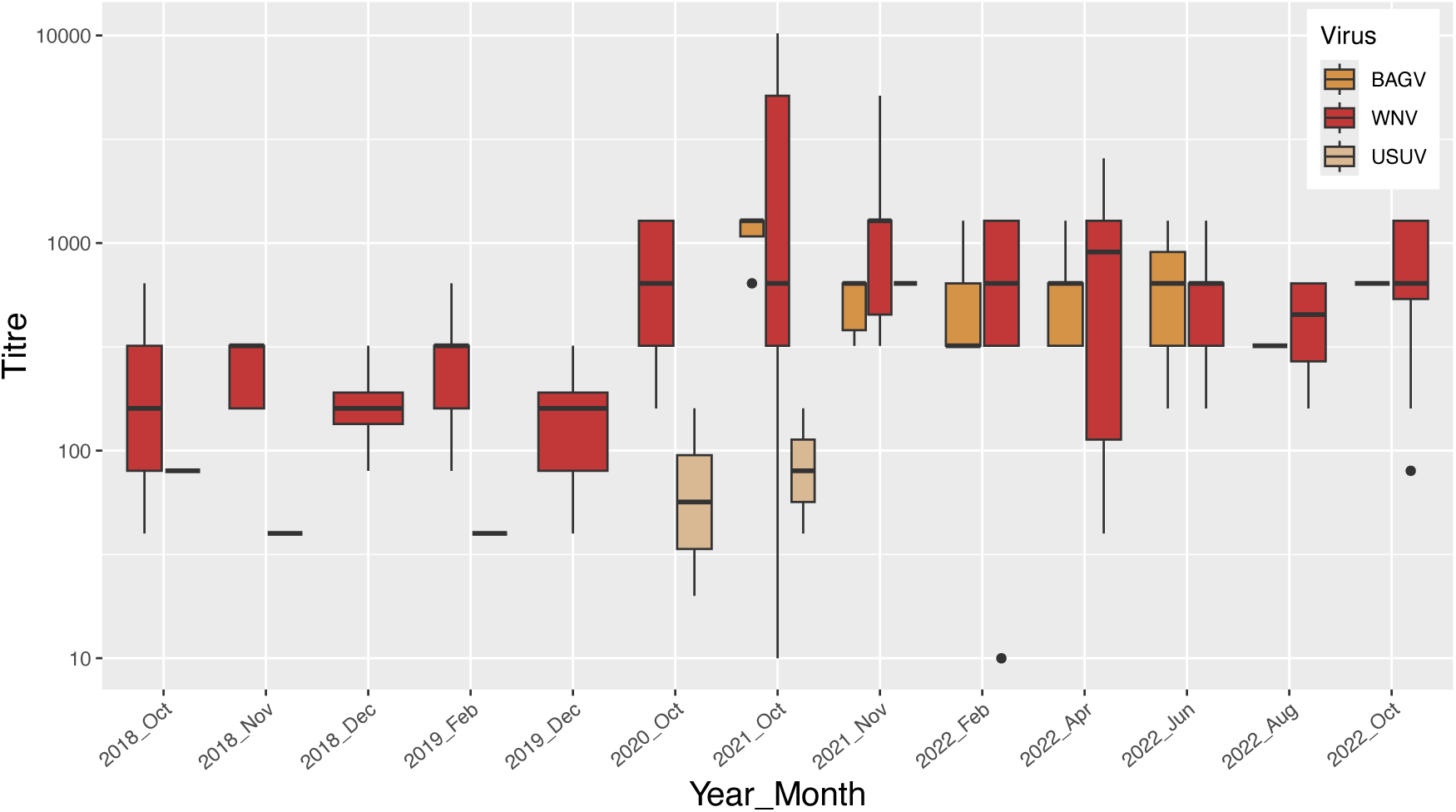
Distribution of the VNT titres to BAGV, WNV and USUV throughout all the sampling seasons. Titres are represented in Log 10 scale. Horizontal dark lines represent the median titre, the boxes represent the interquartile range (25^th^ percentile – 75^th^ percentile), the vertical lines (whiskers) the minimum and maximum values and the dots represent the outliers.

Regarding the drivers of flavivirus exposure, age and year showed significant differences in the ELISA and general VNT models that included the dataset of autumn season across five consecutive years. Seroprevalence was significantly higher in adults in both ELISA (Age Juveniles, β = -1.03, SE = 0.29, p < 0.05) and general VNT (Age Juveniles, β = - 0.96, SE = 0.31, p < 0.05). Concerning the year, for the ELISA results, the years 2020-2022 have a significantly higher flavivirus seroprevalence when compared to 2018 (Year 2020, β = 1.22, SE = 0.54, p < 0.05; Year 2021, β = 0.94, SE = 0.41, p < 0.05; Year 2022, β = 0.93, SE= 0.47, p < 0.05). As for the general flavivirus model, only the year 2021 had a significantly higher flavivirus seroprevalence (β = 1.20, SE = 0.43, p < 0.05).

In the GLM used to compare the drivers of flavivirus seroprevalence after the 2021 BAGV outbreak (Table S5), seroprevalence was significantly higher in males (β = 0.78, SE = 0.37, p < 0.05), except for BAGV. BGAV seroprevalence was significantly lower in August (β = -2.40, SE = 1.16, p < 0.05) and October (β = -2.38, SE = 1.13, p <0.05). No other factors showed significant differences (Table S5).

Although there are no statistically significant differences between adults and juveniles regarding BAGV seroprevalence, a trend can be observed with higher seropositive rates in juveniles (individuals born in 2021) from September/October 2021 to April 2022 (Figure S2). From this date on, the contrary is observed, with higher seropositivity in adults when compared to juveniles born in 2022.

## 4. Discussion and conclusion

Our long-term serological surveillance of red-legged partridges reveals sporadic BAGV and USUV circulation and endemic WNV circulation in Portugal. Understanding the co-circulation dynamics of these viruses in wild birds is a step towards improving surveillance systems at the wildlife-livestock-human interface, particularly for less-known viruses such as BAGV, which has caused recent outbreaks in red-legged partridges in Europe (Queirós et al., 2022; Höfle et al., 2022).

The serological dynamics of BAGV are consistent with its emergence in September 2021 (Queirós et al., 2022), with specific antibodies being detected in September/October 2021, during the first sampling months after the outbreak. In subsequent sampling months, BAGV seroprevalence remains relatively stable until it decreases significantly in August and October of 2022 (Figure 3), when new juveniles are captured and sampled for the first time, suggesting that active circulation ceased or was very low during the 2022 summer season (Figure S2). These results contrast with the serological study carried out in Spain during the 2011-2012 hunting season (October 2011-February 2012), one year after the first Spanish outbreak in September 2010, when the BAGV seroprevalence in game birds (red-legged partridges and pheasants) was 15% (Llorente et al., 2013). Although doubts persist about the duration of BAGV antibodies, our population-level results suggest that antibodies remain detectable in partridges for a limited period of less than one year, which is consistent with the pattern observed for WNV (Pérez-Ramírez et al., 2014).

The serological dynamics of WNV observed in our study are consistent with its endemic circulation in red-legged partridges, with specific antibodies being detected throughout all sampling periods. Although its circulation was expected (Lourenço et al., 2022), the seroprevalences detected in our study (ELISA 58.3% and VNT 28.6%) are higher than those reported previously in Portugal (19.8%, Barros et al., 2011) and in Spain (23%, Llorente et al., 2013; 2.1%-17.5%, Figuerola et al., 2022; 19.23%, Marzal et al., 2022). However, in contrast with previous studies, our seroprevalence data correspond to only one species, highly susceptible to different strains of this virus (Pérez-Ramírez et al., 2018; Sotelo et al., 2011). The prevalence of WNV-specific antibodies was highest in October 2020 (44.8%), just after the large human outbreak detected in southern Spain during the summer of 2020 (Rodríguez-Alarcón et al., 2021).

Taken together, these data indicate that partridges play a relevant role in the WNV transmission/maintenance cycle and suggest the utility of this species as a sentinel for WNV circulation in a given area. The role of other bird species, namely corvids, as sentinels for WNV has already been highlighted (Napp et al., 2019; Tamba et al., 2024). Recently, an integrated surveillance system has shown that active surveillance in corvids allows the detection of WNV before the onset of human cases (Tamba et al., 2024). Corvids, especially magpies, are considered good sentinels, mainly due to their broad distribution and high abundance as well as a high prevalence of WNV antibodies and their sedentarious behaviour and easiness of capture (Napp et al., 2019).

In this study, we detected a high prevalence of WNV antibodies in red-legged partridge, a species that is highly managed in the Iberian Peninsula and is distributed in other European countries such as France, Italy and the United Kingdom, where it was introduced. It is also relatively abundant and easy to capture in some hunting areas and, as a game species, sampling of hunted individuals offers a cost-effective strategy for serological and virological surveillance. Indeed, partridges are susceptible to infection by a wide range of flaviviruses, such as those currently circulating in the south of the Iberian Peninsula (namely, WNV, USUV and BAGV), which makes this species optimal for sentinel monitoring and surveillance of these flaviviruses, a feature that is not clearly established for other bird species suggested for sentinels, singularly corvids.

The serological dynamics of USUV are consistent with sporadic circulation in the red-legged partridge, with specific antibodies being detected in autumn 2018, 2020 and 2021. Seroprevalences were lower when compared with other studies on game birds (red-legged partridges and common pheasants) in Spain in 2011 (10%, Llorente et al., 2013).

Regarding the results obtained for “undetermined flaviviruses”, these could be explained by intrinsic limitations of the laboratory methods, specifically those related to cross-reactions (no specific antibodies to a specific virus are detected even though the infection is caused by one of the three viruses), double infections by more than one flavivirus (either successive or, less likely, simultaneous co-infections) or infection with an unknown flavivirus (Llorente et al., 2019). The “undetermined flaviviruses” seroprevalences seem to follow the tendency for WNV with some influence from other flaviviruses (Figures 2 and 3), namely USUV and BAGV. The increase of undifferentiable reactions to various flaviviruses when both USUV and BAGV first appeared suggests that these “undetermined flaviviruses” could likely be explained by cross-reaction or infection by more than one virus. There is a seroprevalence peak occurring in autumn 2018 in parallel with the first detection of USUV, as well as in September/October 2021 in coincidence with the appearance of BAGV. The fact that the mean titre of the different flaviviruses seems to follow the same tendency also suggests multiple exposure to different flaviviruses.

We detected an overall high flavivirus seroprevalence of 58.1% (274/472) based on ELISA and of 66.3% (285/430) based on VNT. The ELISA test used in this study, although designed for WNV antibody detection, can cross-react with other flaviviruses (Llorente et al., 2019), allowing for a good approximation to the overall flavivirus seroprevalence but not of the specific virus(es) involved. Although VNT is the gold standard technique for flavivirus serological diagnosis, it has some important drawbacks: it has to be performed in a BSL-3 lab, it is very time-consuming and it requires higher amounts of serum, which is not always possible when working with small wild birds. In this context, WNV ELISA tests can be useful as a screening tool for flaviviruses, although they have a strong limitation: WNV antibodies are primarily detected over the rest of flaviviruses, which may vary widely in terms of cross-reactive strength. So, although cross-reactions with other flaviviruses exist, they are less characterized in terms of sensitivity, as compared to WNV, making results difficult to interpret. Since there are no commercial ELISAs available for USUV or BAGV, the VNT is the only tool to discriminate antibodies specific to these two flaviviruses.

Regarding the individual drivers of flavivirus exposure in the autumn, the higher flavivirus seroprevalence in adult partridges in the ELISA and general VNT models might be explained by their long life and consequently higher probability of exposure through mosquito bites. In 2022 seroprevalence was higher in males (except for BAGV). Other studies performed on other bird species have found distinct results, with no differences between males and females (Medrouh et al., 2020) or higher prevalence in females (Egizi et al., 2014), likely explained by higher exposure to mosquito bites during incubation. Since no sex differences in seroprevalence was found in the analyses of the autumn data, these results must be interpreted with caution since sex differences in flavivirus seroconversion are likely an interaction of behavioural, biological and ecological features that are very complex to understand (Egizi et al., 2014).

This is the first study to focus on a long-term flavivirus surveillance system based on a single species in Portugal. The red-legged partridge is a species of high socio-economic importance in the Iberian Peninsula (Farfán, 2022), and it is crucial to deeply understand its role in flavivirus circulation and the impact that BAGV outbreaks may have on its natural populations. This study also shows that continuous monitoring of partridges and other wild birds is important to understand the epidemiological dynamics of BAGV and other zoonotic viruses such as WNV and USUV, highlighting the possible role of these species in the One Health surveillance of emerging viruses in Portugal and Europe, including the identification of risk factors associated with their emergence (BAGV) or persistence (WNV) in wildlife populations.

## Author Contributions

J.Q., U.H. coordinated the study; J.Q, U.H., F.L., E.P-R. designed research; J.Q., U.H., P.C.A, D.G., G.M. acquired funding; J.Q., C.F-G., J.B.C., D.G., P.C.A. contributed with sampling; C.F.-G., F.L., J.Q., U.H. performed laboratory work; C.F.-G., U.H., J.Q. analysed data; C.F.-G, J.Q., U.H., F.L., E. P.-R. M.A.-C. contributed to the interpretation of the results; C.F.-G., J.Q., U.H. wrote the paper; C.F.-G., F.L., E. P.-R., M.A.-C., J.B.C., G.M., D.G., P.C.A., U.H., J.Q. revised the paper and approved the final version.

## Acknowledgements

We would like to thank all the colleagues from Biopolis-CIBIO (Tatiana Silva and other students) and IREC for their kind help in collecting, storing and processing the samples. We are also grateful to Herdade de Vale de Perditos and its staff for their support and help in collecting the samples.

## Ethical statement

This study was carried out in the scope of a surveillance program implemented in a population of red-legged partridge, a common and highly managed game species in the Iberian Peninsula. The programm included capture, marking and collection of biological samples, carried out routinely with several sessions per year. All procedures were carried out by experienced researchers, ensuring the welfare of the animals and with legal permits granted annually by the Portuguese National Institute for the Conservation and Protection of Wildlife (ICNF).

