## Supplementary data for "Dynamics of Bagaza, West Nile, and Usutu viruses in Portugal revealed by long-term serological surveillance in red-legged partridges, 2018-2022"

### Supplementary Tables


Table S1…………………………………………………………………………….2

Table S2…………………………………………………………………………….3

Table S3…………………………………………………………………………….4

Table S4…………………………………………………………………………….5

Table S5…………………………………………………………………………….7


### Supplementary Figures


Figure S1…………………………………………………………………………….9

Figure S2…………………………………………………………………………..10

**Table S1**. Flavivirus seroprevalence calculated using a commercial competitive West-Nile virus ELISA test for the red-legged partridge sampled between 2018 and 2022.

| **Year** | **2018** | | **2019** | | **2020** | **2021** | | **2022** | | | | | **Total** |
|---|---|---|---|---|---|---|---|---|---|---|---|---|---|
| **Season** | **Autumn** | | **Winter** | **Autumn** | **Autumn** | **Autumn** | | **Winter** | **Spring** | **Summer** | | **Autumn** | |
| **Month** | **Oct** | **Nov/Dec** | **Feb** | **Dec** | **Oct** | **Sep/Oct** | **Nov** | **Feb** | **Apr** | **Jun** | **Aug** | **Oct** | |
| **Seroprevalence (%)** | 28.6 (4/14) | 40.9 (18/44) | 62.5 (25/40) | 55.0 (22/40) | 72.5 (29/40) | 66.7 (34/51) | 70.0 (28/40) | 73.3 (33/45) | 64.3 (18/28) | 62.9 (22/35) | 16.3 (7/43) | 68.8 (33/48) | 58.3 (273/468) |
| | 37.9 (22/58) | | | | | 68.1 (62/91) | | | | 37.2 (29/78) | | | |

**Table S2**. Flavivirus seroprevalence calculated using a micro virus neutralization test (VNT) for the red-legged partridge sampled between 2018 and 2022. The number of individuals analysed (n) and the results for each virus (BAGV, WNV, USUV, and "undetermined flaviviruses") are shown.

| **Year** | **2018** | | **2019** | | **2020** | **2021** | | **2022** | | | | | **Total** |
|---|---|---|---|---|---|---|---|---|---|---|---|---|---|
| **Season** | **Autumn** | | **Winter** | **Autumn** | **Autumn** | **Autumn** | | **Winter** | **Spring** | **Summer** | | **Autumn** | |
| **Month** | **Oct** | **Nov/Dec** | **Feb** | **Dec** | **Oct** | **Set/Oct** | **Nov** | **Feb** | **Apr** | **Jun** | **Aug** | **Oct** | |
| **Excluded due to lack of volume (n)** | 3 | 5 | 2 | 1 | 11 | 3 | 2 | 0 | 0 | 3 | 1 | 4 | 35 |
| **Seroprevalence (%)** | | | | | | | | | | | | | |
| **BAGV** | 0 (0/11) | 0 (0/39) | 0 (0/38) | 0 (0/39) | 0 (0/29) | 8.3 (4/48) | 26.3 (10/38) | 15.6 (7/45) | 17.9 (5/28) | 21.9 (7/32) | 2.4 (1/42) | 2.3 (1/44) | 8.1 (35/433) |
| **WNV** | 27.3 (3/11) | 23.1 (9/39) | 36.8 (14/38) | 30.8 (12/39) | 44.8 (13/29) | 27.1 (13/48) | 26.3 (10/38) | 28.9 (13/45) | 35.7 (10/28) | 21.9 (7/32) | 9.5 (4/42) | 36.4 (16/44) | 28.6 (124/433) |
| **USUV** | 9.1 (1/11) | 5.1 (2/39) | 2.6 (1/38) | 0 (0/39) | 6.9 (2/29) | 4.2 (2/48) | 2.6 (1/38) | 0 (0/45) | 0 (0/28) | 0 (0/32) | 0 (0/42) | 0 (0/44) | 2.1 (9/433) |
| **Undetermined flaviviruses** | 9.1 (1/11) | 25.6 (10/39) | 23.7 (9/38) | 20.5 (8/39) | 20.7 (6/29) | 35.4 (17/48) | 31.6 (12/38) | 37.8 (17/45) | 32.1 (9/28) | 34.4 (11/32) | 11.9 (5/42) | 31.8 (14/44) | 27.5 (119/433) |
| **Total** | 45.5 (5/11) | 53.8 (21/39) | 63.2 (24/38) | 51.3 (20/39) | 72.4 (21/29) | 75.0 (36/48) | 86.8 (33/38) | 82.2 (37/45) | 85.7 (24/28) | 78.1 (25/32) | 23.8 (10/42) | 70.5 (31/44) | 66.3 (287/433) |
| | 43.3 (26/60) | | | | | 80.2 (69/86) | | | | 47.3 (35/74) | | | |

**Table S3**. Number of individuals classified as BAGV, WNV, USUV and "undetermined flaviviruses" for each micro virus neutralisation test (VNT) titre category. Titres with "≥" means that no higher dilutions were performed since they were not necessary to differentiate the specific antibodies.

| Titres | BAGV | WNV | USUV | Undetermined flaviviruses | Total |
|---|---|---|---|---|---|
| 1:10 | 0 | 2 | 0 | 13 | 15 |
| 1:20 | 0 | 0 | 1 | 10 | 11 |
| 1:40 | 0 | 3 | 4 | 10 | 17 |
| 1:80 | 0 | 9 | 1 | 10 | 20 |
| 1:160 | 1 | 22 | 2 | 21 | 46 |
| 1:320 | 12 | 32 | 0 | 26 | 70 |
| 1:640 | 15 | 23 | 1 | 13 | 52 |
| 1:1280 | 0 | 3 | 0 | 14 | 17 |
| ≥1:1280 | 7 | 21 | 0 | 0 | 28 |
| 1:2560 | 0 | 3 | 0 | 1 | 4 |
| 1:5120 | 0 | 2 | 0 | 0 | 2 |
| ≥1:5120 | 0 | 1 | 0 | 0 | 1 |
| ≥1:10240 | 0 | 3 | 0 | 1 | 4 |

**Table S4**. Results of the GLMs analysing the relationship between overall flavivirus seroprevalences calculated by ELISA and VNT (General VNT) in the autumn dataset (n = 277). Models were also constructed for the seroprevalence obtained for each virus in the VNT analysis: BAGV, WNV, USUV and "undetermined flaviviruses". Significant results are considered when the *p-value* < 0.05 and are highlighted with bold and *.

| **ELISA ~ Sex + Age +Year** | | | | |
|---|---|---|---|---|
| Coefficients | Estimate | Standard Error | z-value | Pr(>\|z\|) |
| Intercept | 0.015 | 0.393 | 0.038 | 0.9700 |
| Sex Male | 0.385 | 0.283 | 1.360 | 0.1738 |
| **Age Juvenile** | **-1.034** | **0.292** | **-3.538** | **0.0004*** |
| Year 2019 | 0.125 | 0.470 | 0.265 | 0.7908 |
| **Year 2020** | **1.222** | **0.543** | **2.252** | **0.0244*** |
| **Year 2021** | **0.941** | **0.408** | **2.308** | **0.0210*** |
| **Year 2022** | **0.927** | **0.468** | **1.983** | **0.0474*** |
| **General VNT ~ Sex + Age +Year** | | | | |
| Coefficients | Estimate | Standard Error | z-value | Pr(>\|z\|) |
| Intercept | 0.597 | 0.402 | 1.484 | 0.1377 |
| Sex Male | 0.077 | 0.298 | 0.259 | 0.7960 |
| **Age Juvenile** | **-0.962** | **0.308** | **-3.127** | **0.0018*** |
| Year2019 | -0.399 | 0.470 | -0.849 | 0.3959 |
| Year2020 | 0.771 | 0.537 | 1.437 | 0.1508 |
| **Year2021** | **1.197** | **0.429** | **2.793** | **0.0052*** |
| Year2022 | 0.691 | 0.471 | 1.466 | 0.1426 |
| **BAGV ~ Sex + Age +Year** | | | | |
| Coefficients | Estimate | Standard Error | z-value | Pr(>\|z\|) |
| Intercept | -21.340 | 2750.266 | -0.008 | 0.9938 |
| Sex Male | 0.303 | 0.589 | 0.515 | 0.6064 |
| Age Juvenile | 1.076 | 0.584 | 1.843 | 0.0654 |
| Year2019 | 0.274 | 3915.762 | 0.000 | 0.9999 |
| Year2020 | 0.047 | 4243.377 | 0.000 | 1.0000 |
| Year2021 | 19.024 | 2750.266 | 0.007 | 0.9945 |
| Year2022 | 16.805 | 2750.266 | 0.006 | 0.9951 |
| **WNV ~ Sex + Age +Year** | | | | |
| Coefficients | Estimate | Standard Error | z-value | Pr(>\|z\|) |
| **Intercept** | **-0.855** | **0.421** | **-2.033** | **0.0421*** |
| Sex Male | 0.202 | 0.290 | 0.698 | 0.4854 |
| Age Juvenile | -0.572 | 0.307 | -1.863 | 0.0625 |
| Year 2019 | 0.031 | 0.504 | 0.061 | 0.9512 |
| Year 2020 | 0.760 | 0.520 | 1.460 | 0.1442 |
| Year 2021 | -0.070 | 0.434 | -0.163 | 0.8705 |
| Year 2022 | 0.419 | 0.478 | 0.877 | 0.8705 |
| **USUV ~ Sex + Age +Year** | | | | |
| Coefficients | Estimate | Standard Error | z-value | Pr(>\|z\|) |
| **Intercept** | **-2.372** | **0.895** | **-2.652** | **0.0080*** |
| Sex Male | -0.200 | 0.794 | -0.252 | 0.8011 |
| Age Juvenile | -1.520 | 1.100 | -1.381 | 0.1672 |
| Year 2019 | -17.882 | 2795.815 | -0.006 | 0.9949 |

| | | | | |
|---|---|---|---|---|
| Year 2020 | 0.304 | 1.042 | 0.292 | 0.7702 |
| Year 2021 | -0.465 | 0.942 | -0.493 | 0.6219 |
| Year 2022 | -17.622 | 2601.053 | -0.007 | 0.9946 |

| **Undetermined flaviviruses ~ Sex + Age +Year** | | | | |
|---|---|---|---|---|
| Coefficients | Estimate | Standard Error | z-value | Pr(>\|z\|) |
| **Intercept** | **-0.902** | **0.439** | **-2.054** | **0.0400*** |
| Sex Male | -0.195 | 0.297 | -0.657 | 0.5113 |
| Age Juvenile | -0.528 | 0.318 | -1.662 | 0.0965 |
| Year 2019 | -0.247 | 0.556 | -0.444 | 0.6574 |
| Year 2020 | -0.129 | 0.598 | -0.215 | 0.8296 |
| Year 2021 | 0.529 | 0.445 | 1.188 | 0.2347 |
| Year 2022 | 0.461 | 0.502 | 0.918 | 0.3587 |

**Table S5**. Results of the GLMs analysing the relationship between overall seroprevalences calculated by ELISA and VNT (General VNT) in the two-month dataset after the BAGV outbreak (n = 199). Models were also constructed for the seroprevalence obtained for each virus in the VNT analysis: WNV, BAGV and "undetermined flaviviruses". USUV data is not present because no antibodies were detected in 2022. Significant results are considered when the *p-value* < 0.05 and are highlighted with bold and *.

| **ELISA ~ Sex + Age + Month** | | | | |
|---|---|---|---|---|
| Coefficients | Estimate | Standard Error | z value | Pr(>\|z\|) |
| Intercept | -0.950 | 0.486 | -1.956 | 0.0505 |
| **Sex Male** | **0.781** | **0.372** | **2.100** | **0.0357*** |
| Age Juvenile | -0.667 | 0.392 | -1.700 | 0.0892 |
| Month August | -1.394 | 0.761 | -1.832 | 0.0669 |
| Month February | -0.148 | 0.542 | -0.273 | 0.7848 |
| Month June | -0.482 | 0.619 | -0.778 | 0.4367 |
| Month October | 0.216 | 0.533 | 0.404 | 0.6860 |
| **General VNT ~ Sex + Age + Month** | | | | |
| Coefficients | Estimate | Standard Error | z value | Pr(>\|z\|) |
| Intercept | -0.950 | 0.486 | -1.956 | 0.0505 |
| **Sex Male** | **0.781** | **0.372** | **2.100** | **0.0357*** |
| Age Juvenile | -0.667 | 0.392 | -1.700 | 0.0892 |
| Month August | -1.394 | 0.761 | -1.832 | 0.0669 |
| Month February | -0.148 | 0.542 | -0.273 | 0.7848 |
| Month June | -0.482 | 0.619 | -0.778 | 0.4367 |
| Month October | 0.216 | 0.533 | 0.404 | 0.6860 |
| **BAGV ~ Sex + Age + Month** | | | | |
| Coefficients | Estimate | Standard Error | z value | Pr(>\|z\|) |
| **Intercept** | **-1.654** | **0.602** | **-2.748** | **0.0060*** |
| Sex Male | -0.034 | 0.505 | -0.068 | 0.9460 |
| Age Juvenile | 0.568 | 0.533 | 1.065 | 0.2867 |
| **Month August** | **-2.397** | **1.162** | **-2.063** | **0.0391*** |
| Month February | -0.299 | 0.655 | -0.457 | 0.6478 |
| Month June | 0.350 | 0.690 | 0.507 | 0.6123 |
| **Month October** | **-2.372** | **1.134** | **-2.093** | **0.0364*** |
| **WNV ~ Sex + Age + Month** | | | | |
| Coefficients | Estimate | Standard Error | z value | Pr(>\|z\|) |
| Intercept | -0.950 | 0.486 | -1.956 | 0.0505 |
| **Sex Male** | **0.781** | **0.372** | **2.100** | **0.0357*** |
| Age Juvenile | -0.667 | 0.392 | -1.700 | 0.0892 |
| Month August | -1.394 | 0.761 | -1.832 | 0.0669 |
| Month February | -0.148 | 0.542 | -0.273 | 0.7848 |
| Month June | -0.482 | 0.619 | -0.778 | 0.4367 |
| Month October | 0.216 | 0.533 | 0.404 | 0.6860 |
| **Undetermined flaviviruses ~ Sex + Age + Month** | | | | |
| Coefficients | Estimate | Standard Error | z value | Pr(>\|z\|) |
| Intercept | -0.950 | 0.486 | -1.956 | 0.0505 |

| | | | | |
|---|---|---|---|---|
| **Sex Male** | **0.781** | **0.372** | **2.100** | **0.0357*** |
| Age juvenile | -0.667 | 0.392 | -1.700 | 0.0892 |
| Month August | -1.394 | 0.761 | -1.832 | 0.0669 |
| Month February | -0.148 | 0.542 | -0.273 | 0.7848 |
| Month June | -0.482 | 0.619 | -0.778 | 0.4367 |
| Month October | 0.216 | 0.533 | 0.404 | 0.6860 |

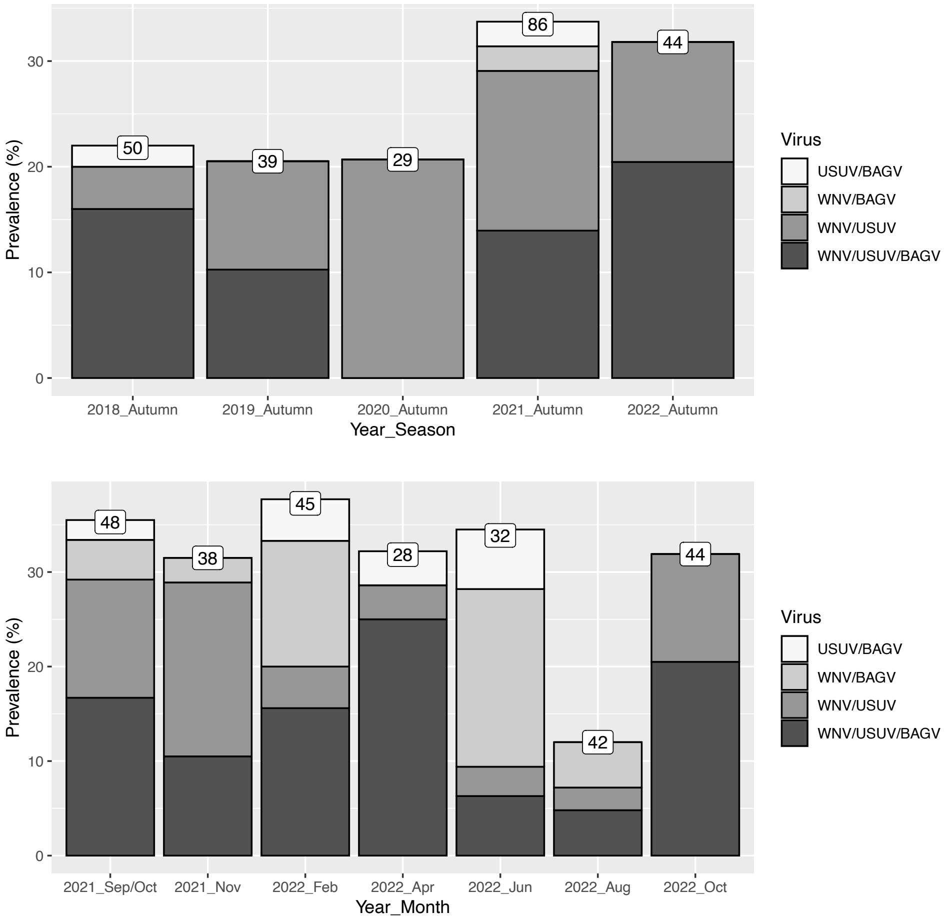


**Figure S1**. Prevalences of the different combinations of virus to which the sera considered "undetermined flaviviruses" reacted to in the micro virus neutralization test (VNT): Above) Throughout the autumn of all years; Below) at two-month sampling intervals after BAGV outbreak. On top of each column is the total number of samples analysed.

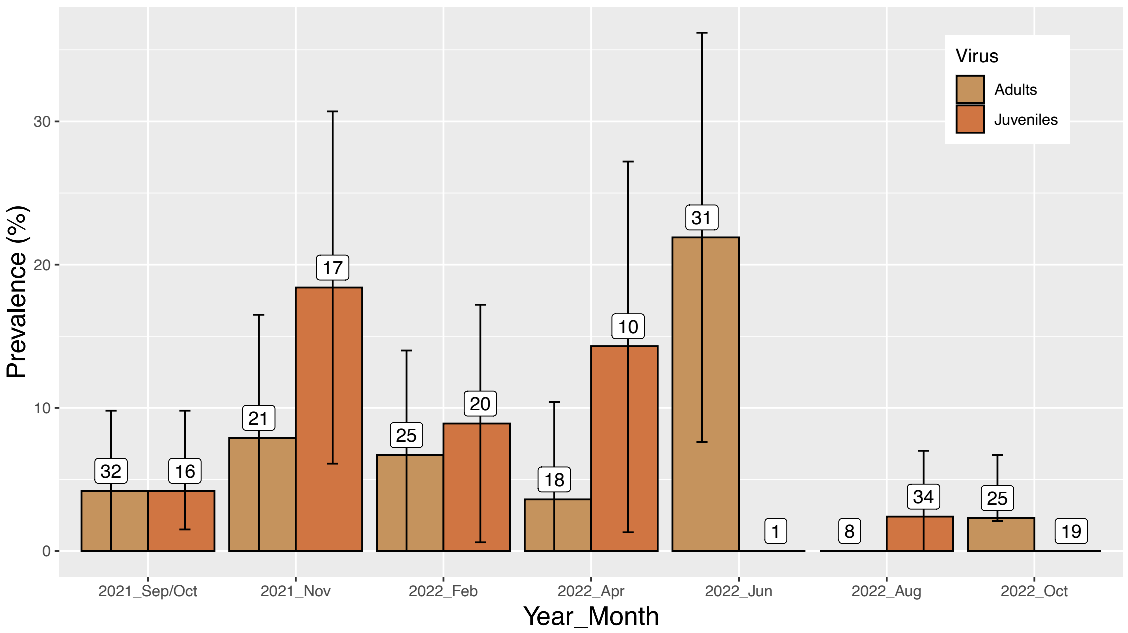


**Figure S2** BAGV seroprevalence obtained by VNT in adults and juveniles after the BAGV outbreak. Individuals born in 2021 are considered juveniles until spring 2022 (April). The number of samples analysed is indicated in each column and the error bars depict the 95% confidence interval.
